# Location, Location, Location: Protein kinase nanoclustering for optimized signalling output

**DOI:** 10.1101/2023.09.29.560079

**Authors:** Rachel S. Gormal, Ramón Martínez-Mármol, Andrew J. Brooks, Frédéric A. Meunier

**Author notes:** Correspondence: Andrew J. Brooks and Frédéric A. Meunier.

## Abstract

Protein kinases (PKs) are proteins at the core of cellular signalling and are thereby responsible for most of the cellular physiological processes and their regulations. As for all cellular proteins, they are subjected to Brownian thermal energy that tends to homogenise their distribution throughout the volume of the cell. To access their substrates and perform their critical functions, PKs localisation is therefore tightly regulated in space and time, relying upon a range of clustering mechanisms. These include post-translational modifications, protein-protein and protein-lipid interactions as well as liquid-liquid phase separation, allowing spatial restriction and ultimately regulating access to their substrates. In this review, we will mainly focus on key mechanisms mediating PK nanoclustering in physiological and pathophysiological processes. We propose that PK nanoclusters act as a cellular unit of signalling output capable of integration and regulation in space and time. We will specifically outline the various super-resolution microscopy approaches currently used to elucidate the mechanisms driving PK nanoscale clustering and explore the pathological consequences of altered kinase clustering in the context of neurodegenerative disorders, inflammation, and cancer.

## Introduction

Protein kinases are an important set of enzymes that play critical cellular roles by controlling cell signalling and a myriad of associated functions. They act by catalysing the phosphorylation of specific target proteins, thereby modulating their activity, subcellular localisation, and function. Through protein phosphorylation, kinases orchestrate essential cellular processes such as cell growth, differentiation, metabolism, and intricate signal transduction. Their mode of action relies on the catalytic transfer of a phosphate group from adenosine triphosphate (ATP) to their selective target proteins (*1*). This phosphorylation acts as a switch in their function by initiating a cascade of molecular events controlling selective cellular behaviours (*2*). Following binding of ATP to the catalytic domain of the kinase, its conversion into adenosine diphosphate (ADP) and inorganic phosphate (Pi) allows the phosphate group (Pi) to be covalently transferred to their protein substrate, leading to the modification of its activity (*3*).

Protein kinases exhibit a high degree of diversity, with various ways to classify them (*4*). One major distinction is into classes based on their specific structural motifs (eukaryotic/atypical) (*5*) and target amino acid residues. The three main classes of protein kinases are serine/threonine protein kinases (STPKs), tyrosine kinases (TKs), and dual specificity protein kinases (DSPKs) (*6*).

*Serine/threonine-specific protein kinases* (STPKs) represent most of the eukaryotic protein kinases. As indicated by their name, they act by phosphorylating serine and threonine residues in their substrate/target protein. These kinases are involved in regulating various cellular processes, including cell growth, proliferation, differentiation, and apoptosis. STPKs play critical roles in signalling pathways, such as the transforming growth factor-beta (TGF-β) pathway and the mitogen-activated protein kinase (MAPK) pathway. They can be further subdivided into ‘classical’ STPKs and atypical STPKs that are only found in certain cell types.

*Tyrosine kinases* (TKs) constitute a distinct class of protein kinases that selectively phosphorylate tyrosine residues. TKs play crucial roles in various forms of cellular communication, proliferation, differentiation, and survival. They can be further categorized into receptor-associated TKs, which participate in receptor signalling pathways, and non-receptor associated TKs that interact with DNA in the nucleus. Prominent examples of tyrosine kinases include the epidermal growth factor receptor (EGFR), the insulin receptor, and the JAK, and Src family kinases.

*Dual specificity protein kinases* (DSPKs) possess the unique ability to phosphorylate both serine/threonine and tyrosine residues. These kinases exhibit versatility in their substrate specificity and often participate in complex cellular processes. DSPKs are involved in regulating cell cycle progression, DNA damage response, and cellular stress signaling.

Protein kinases can also be classified based on their specific functions into signalling protein kinases, metabolic protein kinases, and housekeeping protein kinases. Signalling protein kinases are involved in signal transduction, metabolic protein kinases regulate cellular metabolism, and housekeeping protein kinases perform essential functions within the cell.

The subcellular organisation of PK is of great importance for their signalling function. Our understanding of signalling mainly resolves around a vertical integration of the signal, with ligand binding to receptor, receptor activation and PK initiation of the signaling cascade (Fig.1). With the super-resolution revolution in cell biology, came the realisation that PKs, are organised in clusters with sizes below the diffraction limit of light (*7-9*). The mechanism(s) underpinning such clustering and the functional outcome for PK activity are still open questions, but the emerging concept point to a horizontal integration of the signal revolving around the lateral trapping of receptors and associated PKs in nanoclusters (Fig. 1). Upon binding of the ligand, these nanoscale hubs can in turn generate hubs of PKs and of other downstream effectors raising the possibility that nanoclusters could serve as units of signalling function. Thus, understanding how the signalling output signal is integrated in space and time to generate an output function will require multidisciplinary approaches to solve.

**Figure 1:**
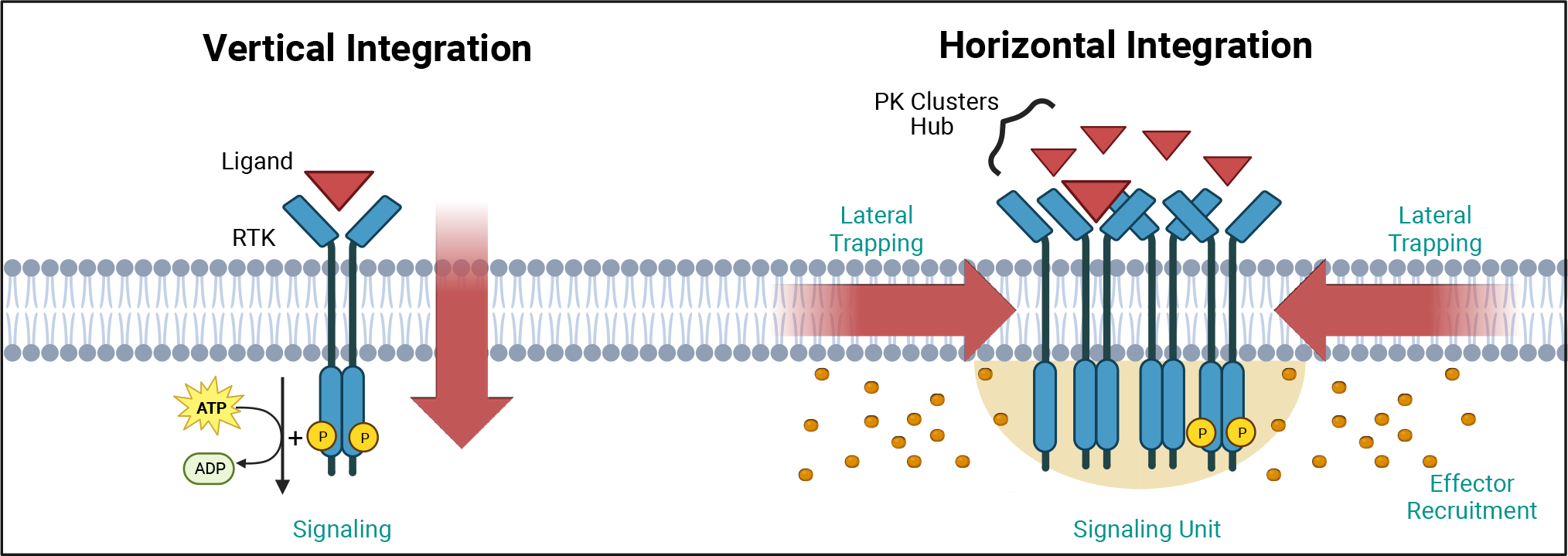
Vertical and horizontal integration of the signal. Protein kinase (PK) activation is classically described as the result of an extracellular ligand (red triangle) binding to a plasma membrane receptor (e.g. RTK) and activating PKs and downstream effectors (orange circles). This vertical integration of the signal has recently been refined with the realisation of the nanocluster organisation of some of the PKs. At the plasma membrane, this horizontal integration occurs via lateral trapping and nanocluster formation of inherently diffusible receptors (and their associated PKs), gives rise to signalling “hubs” or “units”. We have exemplified these two concepts using Receptor Tyrosine Kinases (RTK). Created with BioRender.com.

Due to the large variety of PKs, this review will focus primarily on a limited number of PKs that have been studied with super-resolution microscopy. We will also attempt to draw a roadmap of the use of super-resolution microscopy for assessing the nanoscale organisation of PKs. We will first discuss the main advantages of using sub-diffraction microscopy to assess the dynamics of PK nanoscale organisation, and how this can alter protein signalling cascades in heath and disease. We will therefore briefly cover the main technologies used in these studies. We will then discuss PK membrane clustering, cytoplasmic reorganisation of clustering mediated by lipidation and biomolecular condensates. The review will finally explore the ramifications of PK clustering alterations in neurodegeneration and cancer.

### Super-resolution microscopy for the study of the nanoscale organisation of PKs

Fluorescence microscopy has been instrumental in revealing the subcellular localisation of most PKs, providing additional information on their signalling functions. However, the relatively low resolution of fluorescence microscopy limits our ability to address critical questions inherent to the nanoscale environment in which they operate. The resolution of light microscopy is constrained by the diffraction limit of light which depends on the wavelength of the illumination light used (typically ∼250 nm). Further, many hundreds, if not thousands, of molecules are needed to achieve a signalling output in discrete subcellular locations. In such inherently crowded nano-environments, the fluorescence of so many emitters largely overlap, leading to the detection of large blobby structures lacking resolution. Super-resolution microscopy can, to some extent, overcome the diffraction limit of light. There are 3 commonly used types of super-resolution microscopy techniques (Fig. 2) including stimulated emission depletion (STED) (*10*), structured illumination microscopy (SIM) (*11*) and single-molecule localisation microscopy (SMLM) (*12-14*). SMLM uses the point spread function (PSF) of single fluorescence emitters to fit a Gaussian function and establish the Cartesian coordinates of each emitter with sub-pixel precision, depending on the method used. The PSF is usually circular in the 2 dimensions (x, y) and elliptical in depth (z), and its size depends on the wavelength of the light and the numerical aperture of the objective lens. By fitting a Gaussian function to the PSF, the localisation of the fluorophore can be estimated with relatively high accuracy. There are many excellent reviews that describe the pros- and cons of each super-resolution techniques (*15-17*). One of the key advantages of SMLM is that it can be used both in fixed and live cells (*18*) and is amenable to track endogenous proteins via a range of methods including intrabody expression of selective single-chain antibodies (e.g. camelid nanobodies) (*19*) or CRISPR/Cas9-based endogenous tagging (*20*). They require specialised tags and rely on the activation of a photoswitchable fluorophore that emits sufficient photons for reliable localisation before becoming bleached or going into the dark state (*21*). This is used to track molecules over several consecutive images and derive their diffusion coefficients (*19*). One of the drawbacks of photoactivatable, photoswitchable, and photoconvertible tags is that the trajectories generated are relatively short. Depending on the requirement of the experiments, other techniques based on self-labelling enzymatic tags can be used, such as SNAP tags and Halo tags that are also genetically fused to the target protein. They covalently bind their bright fluorescent cell-permeant ligands, which have excellent photophysical properties (*21*). The sparse labelling obtained with these techniques can be used in fixed and live cells. In the latter case, long trajectories can be generated with high localisation precision. Classically, SMLM techniques have relatively good localisation precision in the range of 10-40 nm (Fig. 2), but new technologies can achieve higher spatiotemporal resolutions (*22, 23*). Overall, a range of super-resolution techniques are well suited to the study of PKs, providing a better understand the relationship between their spatiotemporal localisation and the signalling generated.

**Figure 2.**
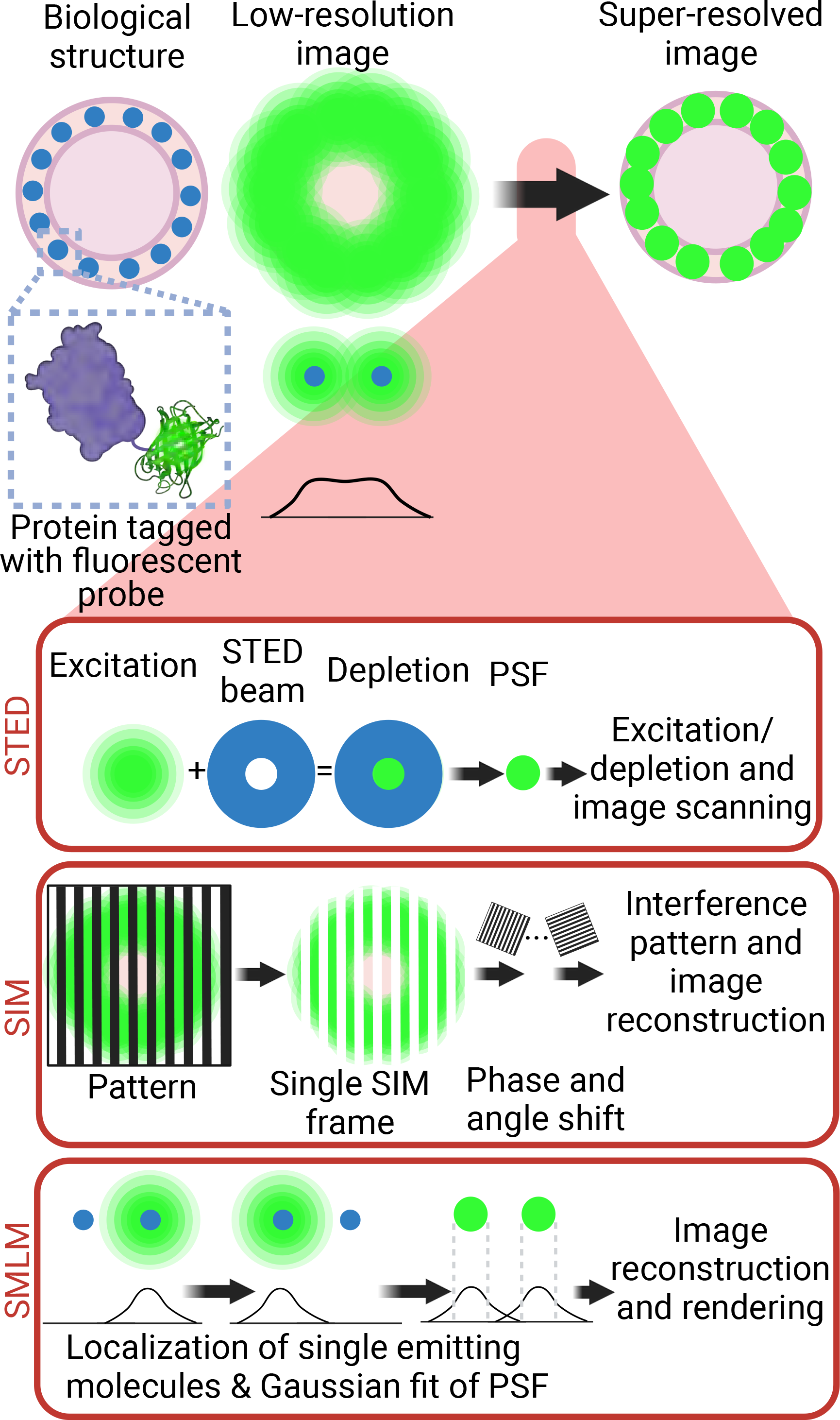
Super-resolution microscopy techniques. Selective labelling of proteins with antibodies or tagging with fluorescent proteins allowed their direct visualization by light microscopy. However, the identification of small cellular structures has been limited by the diffraction limit of light. Only the implementation of super-resolution imaging efficiently solved this limitation. There are 3 main super-resolution approaches. In STED, the focused excitation light is combined with a depletion doughnut-shaped beam, decreasing PSF size to a volume smaller than the diffraction limit. Scanning the sample with an excitation light aligned with STED light beam allows the creation of super-resolved images. In SIM, the sample is imaged with a grid-like pattern of light. The interference patterns between the sample and the illumination grid create Moiré fringes. Multiple images are obtained with varying phase shifts of the patterned light, which are used to reconstruct a sub-diffraction image. In SMLM, the precise position of individual emitting molecules is obtained by fitting their intensity profile to a Gaussian model of the PSF. Acquisition of single localizations depends on the low density and stochastic excitation of emitters. Single localizations are then combined to reconstruct the super-resolved image. Abbreviations: STED, stimulated emission depletion; PSF, point spread function; SIM, structured illumination microscopy; SMLM, single-molecule localization microscopy. Created with BioRender.com.

### Thermodynamic considerations on protein kinases localisation and their signalling output

PK-mediated signalling largely depends on their ability to phosphorylate their substrates. How they access the substrate in a timely fashion is therefore critical to the efficiency and timing of the phosphorylation process. One of the main hindrances to the timely access of PK to substrate comes from Brownian thermal energy, which generates a chaotic nanoscale environment tending at homogenising both PK and their substrates in the entire volume of the cell. The diffusion of an average size protein *in cellulo* is approximately 10 μm^2^/s. This means that cytosolic proteins can navigate the entire length of a cell within a few seconds bouncing against other proteins randomly (*24*). The ability of PKs and their substrates to efficiently react in space and time must therefore require some level of immobilisation. This is a critical factor to take into consideration in the timing and integration of the output signal. In this context, it is not surprising that PK have been found to be immobilised in small subdiffractional clusters, within Receptor Tyrosine Kinases (RTKs) or near (as for Kinase-associated Receptors) the plasma membrane. Nanoclustering of plasma membrane receptors by transient lateral trapping, is emerging as significant for efficient and selective signalling (*25, 26*). The affinity and accessibility of PKs to their target substrates is enhanced by their concentration, and therefore the mechanism(s) controlling the nanoscale organisation of PKs in clusters and that of their substrate is of great interest to the field. In addition, the role of receptor ligand binding on transphosphorylation, hetero-/homo-dimeric/oligomeric re-organisation as well as allosteric competition between downstream effector likely further specialises the downstream response as recently shown (*27*).

Several mechanisms can mediate PK immobilisation influence downstream signalling. This includes the fences and pickets’ plasma membrane model, compartmentalized by actin-based membrane-skeleton “fences” and anchored transmembrane protein “pickets” that are capable of clustering receptor PKs in space and time at the plasma membrane (*28*). Protein nanodomains can also be driven by intra- and inter-molecular interactions including oligomerisation (*29*). Prior to super-resolution approaches, the examination of PK organisation and its effect on cellular signalling has mainly relied upon spatial techniques (*8*) leaving the temporal aspect of clustering unresolved. As such, the contribution of clustering and the role that lateral trapping of cell surface receptor and other PKs plays in their activity, remains to be elucidated.

### Plasma membrane clustering of Receptor tyrosine kinases

Unique to PK-linked receptors is that their intracellular catalytic activity is triggered in response to an external ligand, including a large subset of RTKs (Fig. 3A). Receptor dimerisation or clustering was proposed to be sufficient to initiate the catalytic activity of many RTKs. This is supported by the finding that antibody-mediated clustering (2 epitope IgG binding to 2 receptors) of some RTKs was sufficient to promoter their activation (*30*). However, it is becoming increasingly evident that the precise control of signalling cascades is likely to me multifaceted.

**Figure 3:**
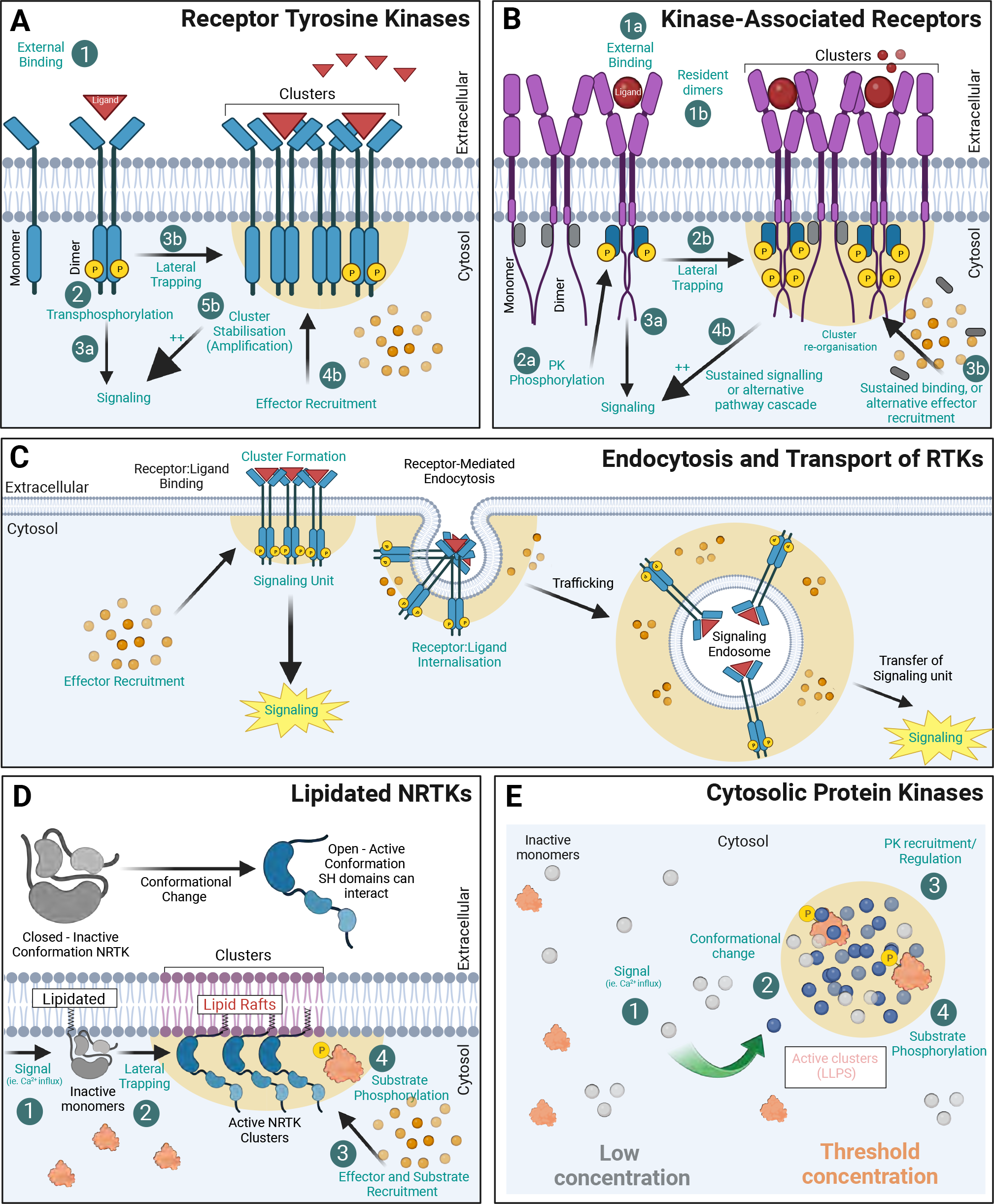
Protein Kinase clustering mechanisms. (**A**) RTKs with intrinsic kinase activity exist at the membrane as both monomers and dimers. Upon ligand binding, trans-phosphorylation of the receptor initiates downstream signalling cascades. The formation of receptor kinase clusters (by lateral trapping) also promotes associated hubs of ligands (red triangles) and effectors (orange circles) that amplify the signal. (**B**) Kinase-Associated Receptors (e.g. cytokine receptors) rely on Non-Receptor PKs to associate to their intracellular domains and activate signalling cascades. Many act through dimerisation (1 ligand:2receptor) and induced conformational change within their intracellular domain (ICD) allowing PK transphosphorylation (grey ‘inactive’, blue ‘active’) and subsequent effector recruitment. (**C**) RTKs undergo receptor mediated endocytosis and endocytic trafficking. Signalling endosomes can continue to signal during transport thereby transferring the “signaling hub” to its subcellular destination. (**D**) Non-Receptor Tyrosine Kinases NRTKs can be post-translationally modified via lipidation (Myristoylated or Palmitoylated) allowing association with membranes. Many of these PKs are allosteric in nature and require a signal (such as Ca^2+^) to alter their conformation and allow subsequent interaction with their substrates. For many, N-terminal myristylation mediates PK association to lipid rafts where they are more active. (**E**) Cytosolic PKs can also compartmentalise into biomolecular condensates (BMCs) which are membraneless compartments, formed via liquid-liquid phase separation (LLPS). Created with BioRender.com.

With the exception of Tyrosine-Kinase associated Receptors, the other four members are able to sense external stimuli through intrinsic catalytic activity/transactivation. Alternatively, TK-associated receptors act through ligand/effector downstream pathways including activation by non-receptor tyrosine kinases (NRTKs). The RTK family of ‘receptor’ kinases is by far the largest superfamily (20 classes) and contains a range of receptors that regulate cell differentiation, proliferation, survival, metabolism, and migration. RTKs utilise several signalling pathways, including MAPK/ERK, PI3K/Akt/mTOR, PLCG1/PKC, JAK/STAT, Ras/Raf, Rac/MEK and NF-kB. Interestingly, RTK signalling occurs in endosomes as well as at the cell surface, with different signalling pathways recruited depending on the location of the receptor. For PK-linked receptors to propagate a signaling cascade, it is thought that several distinct processes must occur: i) ligand binding, ii) activation of the intrinsic kinase domains, and/or iii) binding and activated NRTK to cell signalling (potentially through effectors). Despite the structural differences between TK-Associated Receptors (eg. cytokine receptors) and other intrinsically activated RTKs, the mechanism of ligand binding and re-organisation appears to be somewhat conserved. There are two major, distinct concepts to explain activation of transmembrane, cell-surface receptors. Ligand binding induces either (i) dimerisation of receptors, or (ii) rearrangement of constitutively preformed dimeric receptors.

### Protein kinase cluster by association with plasma membrane receptors

The cytokine receptor Growth Hormone Receptor (GHR) has recently been shown to form nanoclusters at the plasma membrane (*27*) (Fig. 4A). This clustering has been shown to be highly dependent upon competitive binding of two PKs with distinct downstream signalling (*27*). It is unclear whether these clusters are fostering the formation of dimers (*31*), which is needed for effective signalling (Fig. 1B). In the absence of ligand, GHR-ICD is in an inactive conformation. Structural changes in the GHR-ECD induced by the ligand result in an altered ratio of JAK2-STAT5 to ERK1/2 signalling, which has also been shown for other receptors, including the Prolactin Receptor (PRLR) and Erythropoietin receptors (EpoR) (*32-35*). The impact of the competitive binding of PKs for the GHR receptor nanoscale organisation and two alternative signaling pathways was recently demonstrated (*27*). Upon growth hormone (GH) addition and Lyn binding/activation (ERK1/2 pathway), the GHR redistributes into larger clusters on the plasma membrane. Conversely, the activation of the JAK2/STAT pathway did not alter the distribution of the GHR within clusters. A recent study using a single-molecule imaging approach to co-track co-movement of monomeric GHR, and its effectors showed that JAK2 was important in contributing to GHR dimerisation at the membrane (*36*). GHR is able to signal through at least two divergent NTRK pathways, i) JAK2/STAT and ii) LYN/ERK1/2 pathway (*37*). Since the activation of GHR through the Lyn/ERK pathway correlates with increased GHR endocytosis and degradation, its overall membrane organisation may be a key determinant by which a signaling cascade is initiated. In a similar line of research into the cytokine receptor interleukin-2 receptor (IL-2RY) membrane organisation, found that the receptor rarely existed as monomers, and cluster membership increased in response to ligand (IL2) stimulation. In addition, 3 distinct cluster sub-populations have been described: small ‘active’ clusters, medium clusters associated with endocytosis and ‘large’ static clusters (*38*). Perturbation of either lipid rafts or F-actin increased the size of clusters to a similar level, but differentially affected signalling. Cholesterol depletion promoted assembly and sustained STAT5 and ERK signalling, whereas F-actin disruption blocked signalling all together. However, a live cell single-molecule imaging study on the cytokine receptor for thrombopoietin, TpoR, identified a monomer-dimer equilibrium in the absence of Tpo and the ratio of dimers was not significantly altered by Tpo, but that dimers were prolonged. Unfortunately, this study did not extent to investigate clustering (*39*). Another single-molecule tracking study indicated that EGFRs reside outside lipid rafts in the absence of ligand, but move into raft microdomains upon EGF binding (*40*). Lipid raft environment has been shown to enhance LYN kinase activity (*41*). It is therefore likely that the presence of clusters, their size and sub-membrane distribution in regions with different lipidic composition, such as cholesterol-enriched lipid rafts, jointly act to mediate a tight control over the initiation and duration of signalling events.

**Figure 4.**
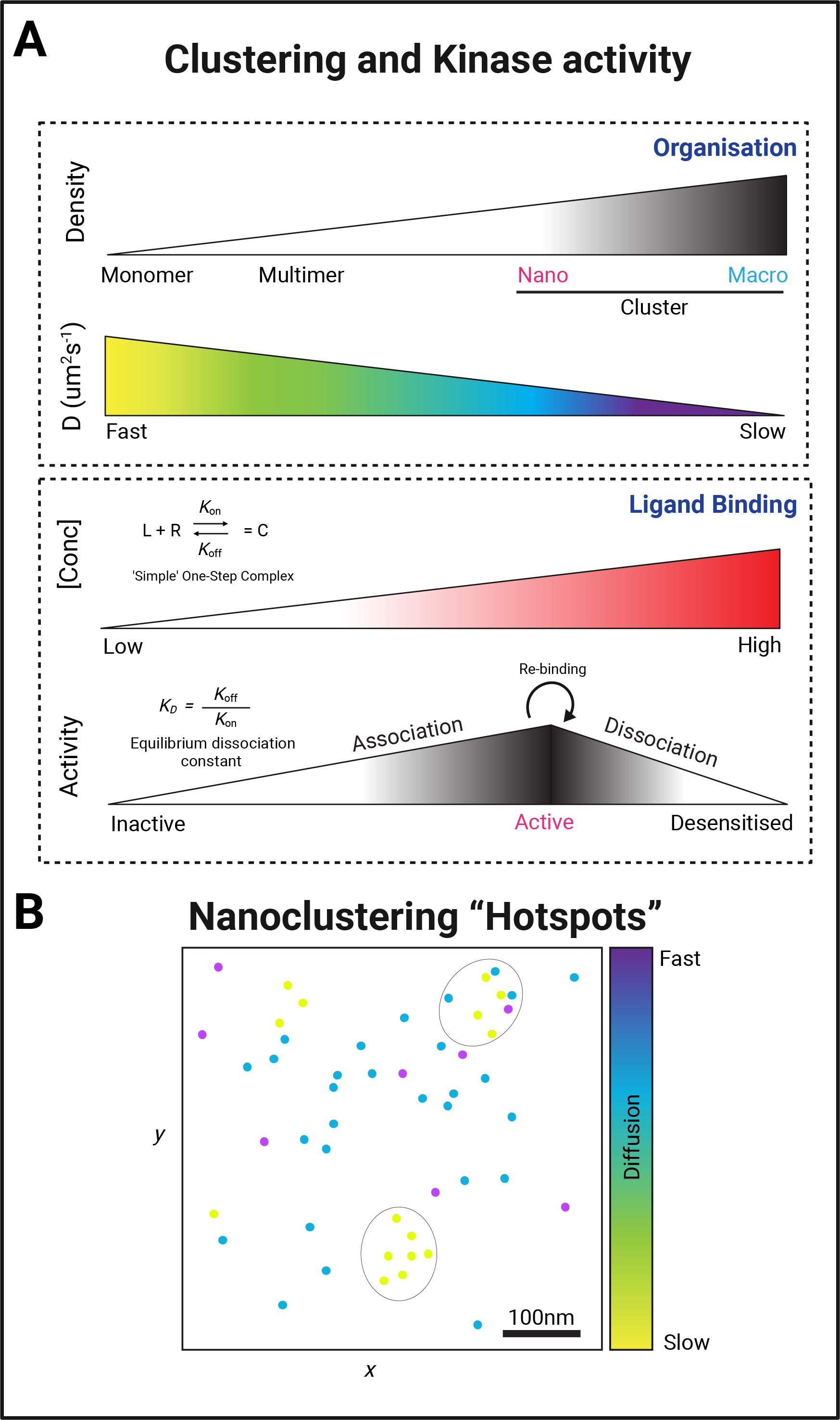
Examples of cellular nanoscale organization of protein kinases. (**A**) Representative sptPALM images of growth hormone receptor tagged with mEos2 fluorescent protein (GHR-ΔBox1(FL)-mEos2) expressed in HEK293 cells. Cells were incubated with human growth hormone (hGH) during the imaging. The panels show the high-resolution intensity map (Intensity), the diffusion coefficient map (Diffusion coefficient), where warmer colors represent lower mobility, and the trajectory map (Trajectories) where warmer tracks appear later in the acquisition. Images modified from (*27*). (**B**) Representative sptPALM images of Fyn-mEos2. The panels show the low-resolution epifluorescence image of a hippocampal neuron expressing Fyn-mEos2 (green) and mCardinal (red) acquired before the photoconversion of mEos2 molecules. The inset is shown at higher magnification in the left panel, where trajectories of single Fyn-mEos2 molecules can be observed. SptPALM imaging was performed as in (*69*). (**C**) Representative sptPALM images of Fyn-mEos2 (left panel) or Fyn-Y531F-mEos2 (right panel) forming nanoclusters in dendritic spines. NASTIC analysis was used for the spatiotemporal distribution of single Fyn-mEos2 trajectories into nanoclusters. Individual trajectories were coloured based on their instant diffusion coefficients (D_eff_), with more immobile trajectories depicted in light colours and more mobile trajectories in dark colours). (**D**) Representative sptPALM images of CamKIIα-mEos2 in neurons. Single CamKIIα-mEos2 localizations were plotted into a diffusion coefficient map (D_eff_) where warmer colors represent higher mobility. The inset shows a dendritic spine at higher magnification in the left panel. Images modified from (*73*).

### Compartmentalisation of PKs in cellular organelles

The endosomal trafficking is critical for a multitude of cellular functions and include transport of various RTKs such as EGFR and platelet-derived growth factor receptor (PDGFR) and PKs associated with receptors such as GHRs (*42*). Importantly, upon cell activation, Src kinase were shown to translocate from the perinuclear region to the plasma membrane on endosomes. An interesting concept is emerging, posing that RTKs-containing endosome could act as signaling units (or ‘quanta’), delivering it to other subcellular destinations (Fig. 3C). This has also been described asas a form of analogue-todigital conversion. This was demonstrated in fibroblasts with EGFR (*43*) and hippocampal neurons with Tyrosine Receptor Kinase B (TrkB) (*44*). The function of this type of endosomalsignaling can differ. For instance, in neurons, axonal retrograde transport of signalling endosomes from the nerve terminal to the soma underpins neuronal survival. Each signalling endosome therefore carries a quantal amount of activated receptors, and it is the frequency of these retrogradely transported endosomes reaching the soma that determines the scale of the neurotrophic signal. Notably, this study demonstrated that pharmacological and genetic inhibition of TrkB activation interfered with the coupling between synaptic activity and retrograde flux of signalling endosomes, suggesting that TrkB activity encodes for the level of synaptic activity experienced distally at nerve terminals and ‘digitalises’ it as flux of retrogradely transported signaling endosomes. The ability of endosomes to signal is not limited to PKs and has also been shown for G-protein coupled receptors (*45*). Therefore, endocytic trafficking of PKs likely represents a general mechanism inherent to the ability of endosomes to generate discrete signalling output (*46*). Further, PKs action is not limited to signaling of the endocytic pathway. Indeed, Src-kinase has been shown to be critically involved in the anterograde pathway by controlling the recruitment of secretory vesicles to the cortical actin network in neurosecretory cells via a direct action and the anchoring property of Myosin VI (*47, 48*).

### Membrane localisation of non-receptor tyrosine kinases via lipidation

One of the most studied mechanisms to control the organisation and function of kinases is by modulating their binding affinity to biological membranes. Non-receptor tyrosine kinases NRTKs are unique, in that their subcellular location is heavily reliant on their conformation, which can be further influenced by a range of post-translational modifications (PTM). Non-receptor tyrosine kinases are categorized into 9 subfamilies based on sequence similarities, primarily within the kinase domains. These includes ABL, FES, JAK, ACK, SYK, TEC, FAK, CSK and Src family of kinases (SFKs). The Src family kinases (SFKs), are active mediators of signal transduction pathways that influence cell proliferation, differentiation, apoptosis, migration, and metabolism. The SFKs members include: Src, Lck, Hck, Blk, Fgr, Lyn, and Yrk. SFKs all contain modular Src homology domains (SH2, SH3, SH4) at their N-terminus, a tyrosine kinase domain (SH1) and intrinsically disordered region (IDR) at its C-terminus (Fig. 3D). The intramolecular interaction of SFKs SH domains has been shown to mediate two main protein conformations, (i) open and (ii) closed (when interacting). The PTMs identified to influence these cytoplasmic receptors include ubiquitination, phosphorylation and lipidation (*49*). Lipidation is essential to facilitate membrane attachment to those peripheral kinases that do not contain membrane-spanning domains. Lipidation consists of the covalent binding of specific lipid moieties to the protein body. Kinase lipidation can be performed co- or post-translationally, where at least two types of lipids have been described attached to kinases, including different forms of fatty acids and lipid-derived electrophiles (LDEs).

Protein kinase localisation and biological function can be regulated by the addition of two types of saturated fatty acyl chain, the 14-carbon myristic acid (C14:0) or the 16-carbon palmitic acid (C16:0). Depending on the type of lipid incorporated, the fatty acylation can be classified as N-myristoylation or S-palmitoylation respectively.

*N*-Myristoylation is an irreversible co-translational protein modification catalyzed by *N*-myristoyl transferases (NMTs), where the fatty acid myristate is covalently attached to the N-terminal glycine of a protein. Both human NMT isozymes, NMT1 and NMT2, are expressed in most tissues and have been implicated in the development and progression of diseases including cancer (*50*), epilepsy (*51*), Alzheimer’s disease, (*52*) and Noonan-like syndrome (*53*). Examples of myristoylated kinases and their function can be found in the Table 1, being SFKs some of the most studied lipid-modified kinases. In fact, all members of the SFK family are co-translationally myristoylated at the second glycine (*54*). Myristoylation is essential to anchor kinases in the cytoplasmic face of the plasma membrane and regulate their enzymatic activities, with important consequences for the organism. High-fat diet favours myristoylation and overactivation of the c-Src kinase, which accelerates xenografted prostate tumour growth in mice *in vivo* (*55*). Myristoylation also regulates kinase protein levels. Mutant c-Src lacking myristoylation showed reduced kinase activity but had enhanced stability, as degradation by ubiquitination was diminished. This effect could be associated with the ability of myristate to facilitate the targeting of c-Src to the membrane, or to an intrinsic requirement for the myristate lipid in promoting ubiquitination and degradation by the E3 ligase Cbl (*56*). Whereas myristoylation positively regulates most c-Src kinase activity, there are exceptions, such as the blockade of c-Abl tyrosine kinase by addition of a myristoyl group. Under resting conditions, c-Abl is in an inactive state maintained by the binding of the N-terminal myristoyl group to a hydrophobic pocket in the C-lobe of the kinase domain (*57*).

**Table 1:**
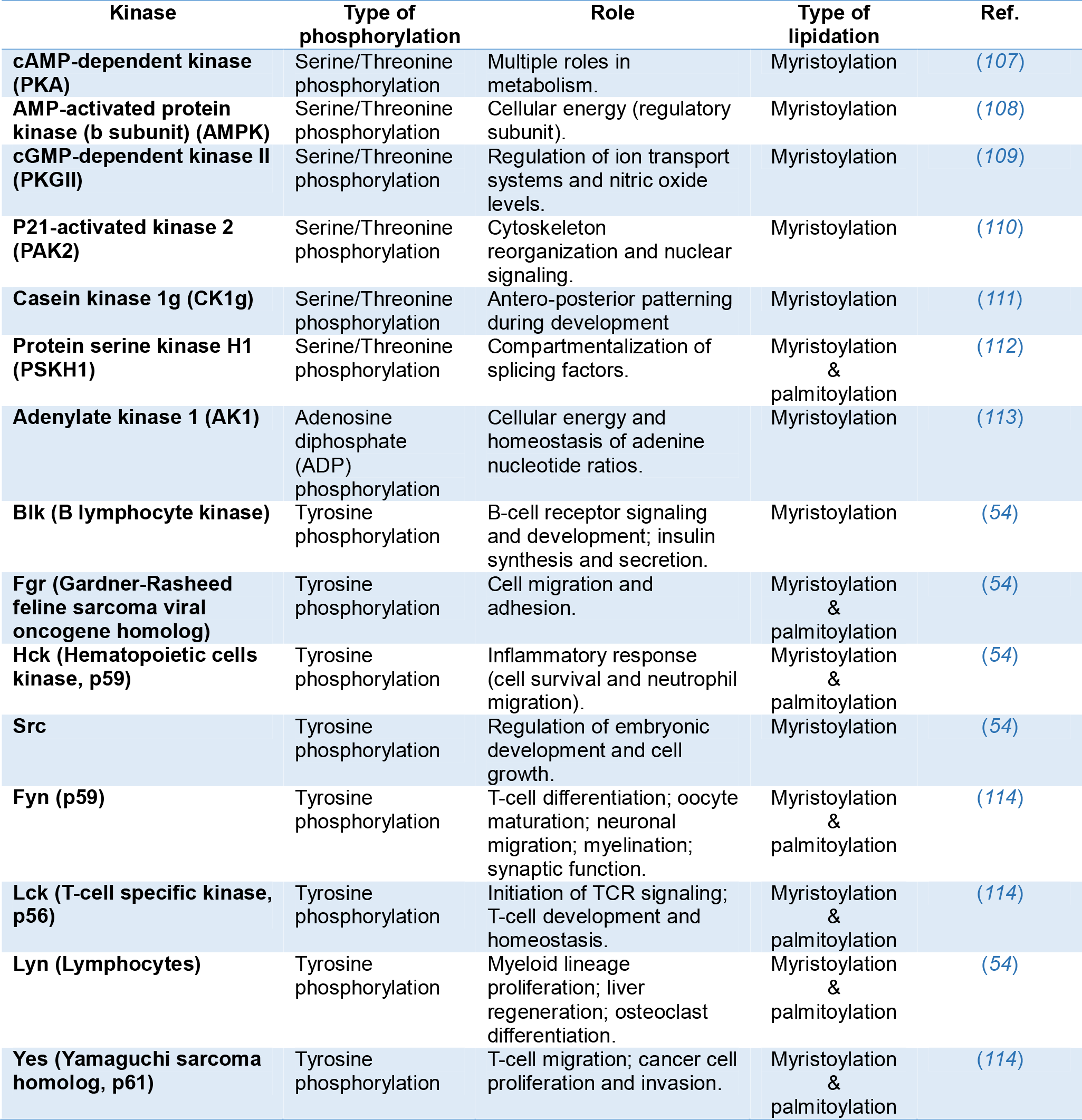
Lipidation types of Protein Kinases.

*S*-palmitoylation (or *S*-acylation) is another type of fatty acylation, in which the fatty acid palmitate is covalently attached to cysteine residues. Contrary to myristoylation, palmitoylation of proteins happens post-translationally and is a reversible process catalysed by a large family of proteins known as protein acyl transferases (DHHC-PATs) (*58*). The labile nature of the thioester bond makes palmitoylation a reversible and very dynamic process, where deacylation is performed by acyl protein thioesterases (APTs) or the lysosomal palmitoyl protein thioesterases (PPTs). The regulated activity of DHHC-PATs and APTs or PPTs facilitates a rapid turnover of membrane-bound palmitoylated proteins that can undergo cycles of acylation and deacylation in response to upstream signals (*59*). Palmitoylation can also modulate the trafficking of the kinases. Mutant forms of Fyn lacking two palmitoylatable cysteines (cysteine 3 and 6) showed altered neuronal distribution, being unable to reach the dendritic arbour (*60*). Palmitoylation is usually found in combination with other lipid modifications. Several SFKs, including Fyn, Lyn, Lck and Yes undergo both myristoylation and palmitoylation (Table 1). Whereas myristoylation alone facilitates the target of proteins to discrete membrane compartments, plasmalemma association of sole-myristoylated proteins is only transient with very short half-lives. The combination with palmitoylation mediates a stronger and longer membrane association. A peptide probe comprising the amino-terminal myristoylated and palmitoylated heptapeptide of the Fyn kinase showed an anterograde transport with initial Golgi accumulation before reaching the plasma membrane. Whereas palmitoylation was detected only on the Golgi, accelerating the anterograde transport of acylated targets, depalmitoylation occurred everywhere in the cell. The plasma membrane localisation of the peptide probe at steady state was more pronounced than other probes due to lower palmitate turnover kinetics (*61*). Similarly, newly synthesized myristoylated Lyn and Yes initially enter the Golgi system, where they become palmitoylated, providing necessary access to the membrane secretory transport pathway *en route* to the plasma membrane, where Rab11 is involved (*62*). Deletion of three base pairs in the *Zdhhc21* gene resulted in the “depilated” phenotype (*dep*), characterised by a variable hair loss, with thinner and shorter hairs remaining. This single mutation resulted in the loss of a highly conserved phenylalanine at position 233, causing the mislocalisation and loss of catalytic activity of the ZDHHC21 acyl transferase. Reintroducing WT ZDHHC21 into the mice rescued the shiny and smooth coat phenotypes. Fyn is a substrate for ZDHHC21, and the lack of palmitoylation caused altered Fyn localisation and downstream signalling activity, that resulted in reduced levels of Lef1, nuclear β-catenin, and Foxn1, altering keratinocyte differentiation that led to hair loss (*63*). Overall, these results indicate that modulation of de-*A*cylation/re-*A*cylation cycles is an important mechanism that control spatially and temporally the activity of SFKs, hence regulating a multitude of downstream signalling cascades with direct consequences *in vivo*.

*Other types of lipidation can also occur such as lipid-derived electrophile (LDE) modification of the ZAK kinase*. LDEs are reactive lipid metabolites generated by lipid peroxidation when cells are under oxidative stress conditions. One important example of LDE is 4-hydroxy-2-nonenal (4-HNE), which is formed as a secondary intermediate by-product of lipid peroxidation (*64*). 4-HNE is a reactive molecule that participates in multiple physiological processes as a nonclassical secondary messenger and can be covalently attached to numerous proteins, including the ZAK kinase (*65*). This kinase is a member of the mitogen-activated protein kinase kinase kinase (MAP3K) family of signal transduction proteins, activating all three major MAPK pathways in mammalian cells (ERK/2, JNK1/2/3 and p38 MAPK) (*66*). ZAK kinase is involved in the cellular response to UV radiation (*67*) and to chemotherapeutic agents (*68*). 4-HNE binds to ZAK in the conserved cysteine 22. The proximity of this position to the ATP-binding loop of ZAP, resulted in a 4-HNE-dependent blockade of ATP binding and loss of kinase activity. This creates a negative feedback mechanism that suppresses the activation of JNK apoptotic pathways induced by oxidative stress.

### Compartmentalisation of PKs in biomolecular condensates

Cytosolic PKs can also be organised in small clusters occurring outside the context of the plasma membrane or organelle membranes(*69*). These sub-diffractional structures have commonly been defined as biomolecular condensates (BMCs) that are generated through liquid-liquid phase separation (LLPS) have been commonly named membraneless organelles (Fig. 3E). These subcellular structures are defined by their ability to concentrate proteins and other charged and disordered molecules such as nucleic acids (*70*). This compartmentalisation creates a unique environment for proteins and is mediated by weak interactions, mostly via unstructured region of proteins also known as intrinsically disordered regions (IDR). Since, the probability of binding between two interacting molecules increases with the square of the binding density (*71*), BMCs are ideally positioned to host and/or initiate highly efficient cellular signalling. These phase-separated condensates are increasingly viewed as critical in a multitude of biological contexts. BMCs can also interface with the plasma membrane and generate hybrid systems, formed through the interplay of phase-separated condensates and membrane constituents, such as transmembrane surface receptors. The Src-kinase Fyn was recently shown to form nanoclusters in dendritic spines (Fig. 4B, B) that were controlled by the ability of Tau to form BMCs (*69*). As Fyn kinase binds to a number of post-synaptic proteins critically involved in synaptic plasticity, it is tempting to speculate that the integration of the signaling and plastic response could be regulated by nanoscale BMCs. Further protein phosphorylation and dephosphorylation can in turn modulate the fabric of these condensates thereby generating multi-layered signal regulation. A recent study mapped a large number of phosphosites enriched within purified condensates, finding phosphosites modulating protein–RNA interactions and impacting their ability to populate their condensates (*69*). Ca^2+^/calmodulin-dependent protein kinase II (CaMKII), is a postsynaptic kinase that acts as a protein cross-linker, segregating synaptic molecules through Ca^2+^-dependent LLPS formation (*72*). CaMKII organises forming nanodomains in dendritic spines (Fig. 4D) (*73*). These nanodomains are essential for the establishment of trans-synaptic nanocolumns (*74*), which may be involved in stablishing an optimal spatial arrangement between postsynaptic receptors and the location where neurotransmitters are released, a key mechanism for neuronal communication and synaptic plasticity. The composition, structure, formation, and role in signaling of BMCs are the subject of intense scrutiny. It is interesting to note that most *in cellulo* work involves large BMC structures amenable to fluorescent recovery after photobleaching. Classically, recovery from photobleach BMC is much slower than from the cytosol. In most cases, these studies have been performed in the context of protein overexpression. Whether kinase residency within BMCs is physiologically relevant is under debate. Recent studies suggest that BMCs can affect the clustering of PKs in the nanoscale range (*69*).

### The role of protein kinases altered clustering in disease

Dysregulated receptor and kinase signaling is a common mechanism driving cancer progression (*75, 76*). Both cholesterol content and modification of phosphoinositide lipids affect transmembrane receptor clustering and signalling (*38, 77-82*). Clinical observations provide support for the importance of cholesterol in regulating cytokine receptor signalling as hypercholesterolemia is associated with leukaemia and has been shown to amplify cytokine signalling in leukaemia cells and alter SFK activation (*83, 84*). In addition, Chronic Lymphocytic Leukaemia (CLL) patients show a survival benefit from cholesterol lowering statin drugs (*85*). Cholesterol levels have also been implicated in regulating other diseases for which cytokine signalling plays a major role, such as rheumatoid arthritis, where elevated LDL cholesterolaemia correlates with increased disease progression.Treatment with statins show anti-inflammatory effects and reduced disease symptoms (*86, 87*). Depletion of membrane cholesterol (which inhibits lipid rafts) inhibits JAK activation by ligand binding to GHR and IL-7R (*88, 89*).

Interactions of receptors with phosphatidylinositol-4,5-bisphosphate (PtdIns(4,5)*P*_2_) have shown to be an important mediator of receptor clustering and signalling. For example, EGFR forms significantly larger and more abundant nanoclusters in the membrane of lung cancer cells compared to normal lung epithelial cells. EGFR clusters are mediated by interaction with PtdIns(4,5)*P*_2_, and PtdIns(4,5)*P*_2_ depletion disrupts EGFR plasma membrane clustering, impairing EGFR signalling. Residues in the EGFR juxtamembrane (JM) region mediate the interaction with PtdIns(4,5)*P*_2_ and mutation of these JM residues similarly disrupts clustering and EGFR signalling (*78*). The cytokine receptors GHR and PRLR, as well as the associated JAK2, are known to have specific interactions with PtdIns(4,5)*P*_2_. Mutations of residues in prolactin receptor (PRLR) that interact with PtdIns(4,5)*P*_2_ impair receptor signalling (*90, 91*).Further, Ras (small GTPase) nanoclustering have been shown to be important in oncogenic signalling and attempts to disrupt Ras nanoclusters are being explored as a potential therapeutic strategy (*92*). Several of studies have identified Ras interaction with phosphatidylserine in membrane clustering and a recent super-resolution microscopy study defined the nanoscopic spatial association in the membrane between Ras and Phosphatidylserine (*93*).

Src family kinases (SFKs) are aberrantly activated in cancer, particularly in solid tumours (*94*). SFKs have been shown to play important roles in the clustering and mobility of receptors such as nicotinic acetylcholine receptors, GPI-anchored receptors, B-cell receptors, and cytokine receptors (*27, 95-98*). Increased membrane-saturated fatty acids induce c-Src clustering within membrane subdomains and activation. This has been postulated as a mechanism contributing to increased type 2 diabetes with obesity (*99*). Increased membrane cholesterol suppresses the transforming ability of SFKs in fibroblasts by being distributed to the cholesterol-enriched membrane microdomains that sequesters them away from signalling pathways (*83*). Proliferation of myeloproliferative neoplasms is commonly due to constitutive signalling of the cytokine receptor TpoR (MPL)(*82*).

Since Protein kinases play a major role in so many biological functions, alterations in these proteins affecting conformation and PTM play a particularly significant role in the disease progression of cancer, infectious diseases, and neurological disorders. The dysregulation of RTKs has been proposed for almost all forms of human cancer since they play such a major role in cell division. The tight regulation of these catalytically active receptors is key to regulating unwarranted cell division and tumorigenesis (*100*). In addition, the JAK-STAT signalling pathway is an intrinsic driver or metastasis (*75*).

### PK nanoclustering for optimal output signal

The binding affinity of ligands to their targets, such as a ligand to a cell surface receptor, or a protein kinase to its substrate is an important metric that ultimately controls signalling strength and duration. However, there are other factors that can make a substantial contribution to the signal output, such as local concentrations, protein conformational states as well as molecular crowding. The terms of affinity and avidity are often used in the context of antibody binding, but the principles of avidity/multivalency have also been applied to characterise several other types of interactions (*101*), such as intermolecular interactions (allosteric proteins and IDPs) (*102*), and interactions associated with short linear motifs (SLiMs) (*103*). Since receptors and associated PKs can potentially participate in more than one signaling cascade, the mechanism by which they can favour one over the other is still mostly elusive. Other factors in determining the nature and strength of the output signal are (i) the probability of ligand/binding, activation and dissociation (*104*), (ii) their multimeric state (*27, 31*), (iii) their diffusive and clustering properties (*105*), the local concentration (*106*) (Fig. 5). It has therefore been proposed that the residence of proteins within sub-cellular compartments, such as lipid rafts or BMCs, likely serves to modulate their activity through dynamic clustering mechanisms (*101*). More experimental and *in silico* work are needed to elucidate the precise cellular organisation of protein kinases to optimally amplify and locally control signaling. The ability for a single type of receptor to activate numerous signaling responses likely relies upon these biophysical properties to modulate Receptor Tyrosine Kinases, Kinase-Associated Receptors as well as Non-Receptor Protein kinases.

**Figure 5.**
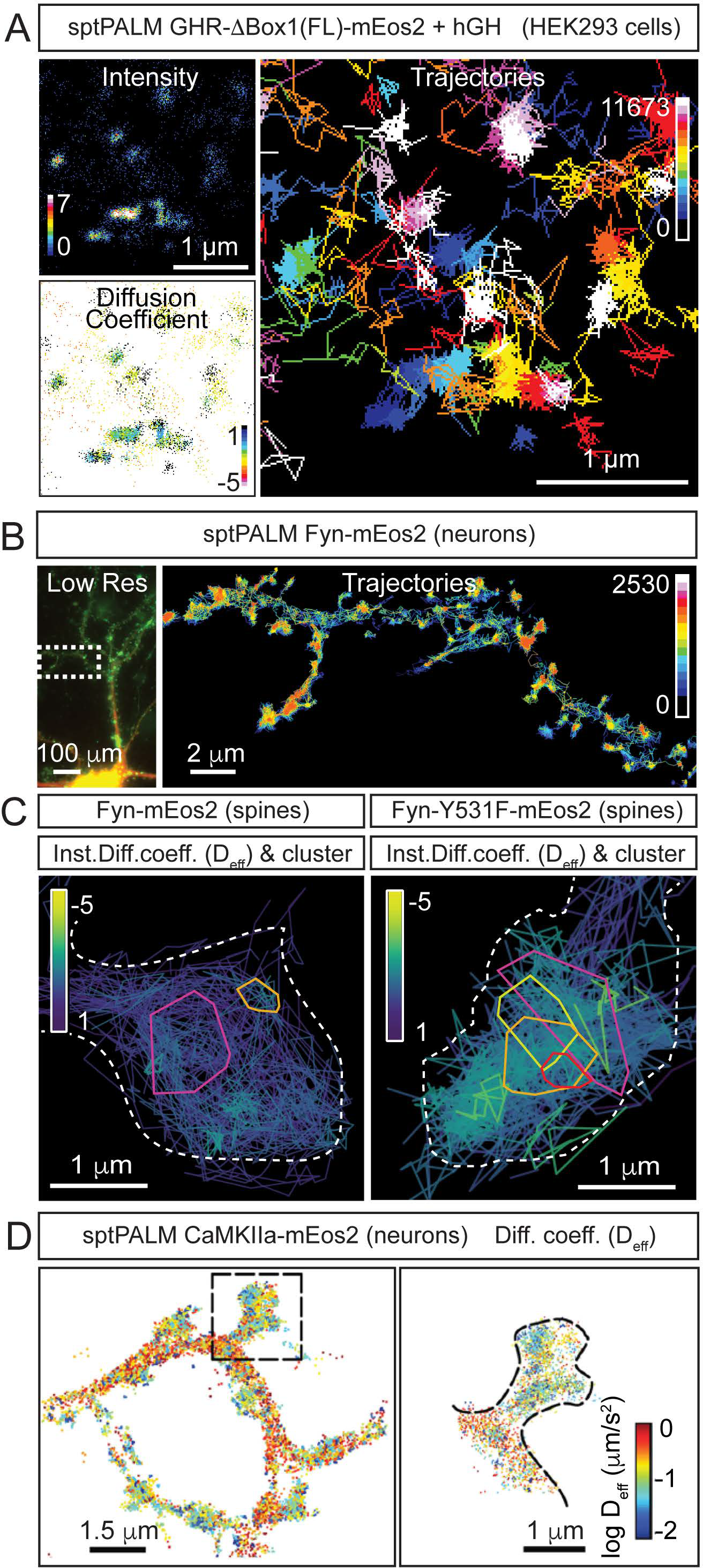
Forming protein kinase clusters. (**A**) PK clustering occurs via lateral trapping in nanoclusters that restrict their diffusion. In addition, the concentration of both ligand and receptor/substrate availability and the dissociation metrics of their interaction have been described to play a key role in determining the signalling strength of this biological event. (**B**) The regulation of both the size and location of clusters is therefore likely to determine the signalling duration through the creation of a nano-environment (circular areas containing a higher density of slow diffusing PKs) conducive for fast re-binding of proteins with their substrates and regulated signalling amplification through the creation of hubs that allow efficient effector association.

## Acknowledgments

F.A.M is supported by a National Health and Medical Research Council (NHMRC) Fellowship (GNT1155794) and an Australian Research Council Discovery Project (DP190100674). The work was supported by UQ Research Stimulus Allocation 2 (postdoctoral fellowship to R.M.M) and two NHMRC grants (APP1157348 and APP1084797) to A.J.B.

